# Dopamine regulates the membrane potential and glycine release of AII amacrine cells via D1-like receptor modulation of gap junction coupling

**DOI:** 10.1101/2024.12.11.625486

**Authors:** Paulo Strazza Junior, Colin M Wakeham, Henrique von Gersdorff

## Abstract

Dopamine plays a pivotal role in adjusting the flow of information across the retina as luminance changes from night to day. Here we show, under dim photopic conditions, that both dopamine and a D1-like receptor (D1R) agonist hyperpolarized the resting membrane potential (V_m_) of AII amacrine cells (AII-ACs). Surprisingly, in the presence of glutamatergic and GABAergic synaptic blockers that isolate glycinergic synapses, D1R agonists are without effect. However, a D1R antagonist depolarized V_m_ and reduced the input resistance of AII-ACs in wild type mice, but not in *Cx36^−/−^* mice. Accordingly, D1R antagonists enhanced tonic glycinergic transmission to type-2 OFF-cone bipolar cells (OFF-CBCs). D1Rs thus adjust the V_m_ and excitability of AII-ACs and, thereby, the level of glycine release to OFF-CBCs by regulating gap junction coupling with ON cone bipolar cells. Our findings provide insights into how the retina may use dopamine to adapt crossover inhibitory microcircuits during changes in luminance.

## Introduction

AII amacrine cells (AII-ACs) are glycinergic interneurons of the mammalian retina that play a crucial role in both scotopic (night) and photopic (daytime) vision (Manookin, Beaudoin, Ernst, Flagel, & Demb, 2008; Strettoi, Masri, & Grünert, 2018). These cells are primarily responsible for converting excitatory signals from rod bipolar cells (RBCs) and ON-cone bipolar cells (ON-CBCs) into glycinergic inhibition of OFF-cone bipolar cells (OFF-CBCs) and are therefore regarded as central hubs of retinal crossover inhibition (Marc, Anderson, Jones, Sigulinsky, & Lauritzen, 2014). The distal (arboreal) dendrites of AII-ACs receive glutamatergic inputs from RBCs and gap junction inputs from ON-CBCs, and these signals then cross to specialized lobular appendages on their proximal (root) dendrites where they can elicit glycine release from a large pool of synaptic vesicles (Demb & Singer, 2012; Mills & Massey, 1995; Strettoi, Raviola, & Dacheux, 1992; Wässle, 2004). Crossover inhibition enhances the signal-to-noise ratio and contrast encoding of retinal circuits (Liang & Freed, 2012). In addition to reciprocal synapses, AII-ACs are also coupled to each other through homotypic connexin 36 (Cx36) gap junctions. The coupling between AII-ACs and ON-CBCs can be either homotypic or heterotypic (Cx36-Cx45) and is modulated by D1 receptors (Bloomfield & Völgyi, 2009; Bloomfield, Xin, & Osborne, 1997). AII-ACs are also unique in the retina in their ability to fire small amplitude Na^+^ spikelets that have been proposed to encode and accelerate signals from RBCs (Tamalu & Watanabe, 2007; Tian, Jarsky, Murphy, Rieke, & Singer, 2010).

AII-ACs are extensively contacted by dopaminergic amacrine cells (DA-ACs) which form ring-like appendages around their somas and other *en passant* synapses around their proximal dendrites (Debertin et al., 2015; Voigt & Wassle, 1987). Through a dense plexus of varicosities and dendritic projections that extend all over the outermost region of the inner plexiform layer (IPL), DA-ACs also make contacts with several other amacrine cell subtypes (e.g., A8 and A17). Some processes also extend to deeper regions of the IPL or to the outer plexiform layer (OPL), where synaptic contacts can form with OFF-CBCs, ON-CBCs, intrinsically photosensitive retinal ganglion cells (ipRGCs), horizontal cells, and cone photoreceptors (Kolb, 1991; Roy & Field, 2019). Virtually all major cell types in the retina, from photoreceptors to ipRGCs, express one or more types of dopamine receptors (Roy & Field, 2019; Warwick, Heukamp, Riccitelli, & Rivlin-Etzion, 2023), which are classified into two families: D1-like (D1 and D5 receptors) and D2-like (D2, D3, and D4 receptors). Typically, D1-like receptors (D1Rs) are coupled to G_αs_- and/or G_αq_-proteins, while D2-like receptors are coupled to G_αi_- and/or G_βg_-proteins (Beaulieu, Espinoza, & Gainetdinov, 2015).

Dopamine release can be synaptic and by volume transmission (paracrine) and is influenced by ambient light levels (Bjelke et al., 1996; Puopolo, Hochstetler, Gustincich, Wightman, & Raviola, 2001). Dopamine release is thought to be very high in bright light conditions, reduces to low basal levels under dim light or twilight, and increases slightly in dark conditions (Gustincich, Feigenspan, Wu, Koopman, & Raviola, 1997; Marshak, 2001; Pérez-Fernández et al., 2019). Dopamine is known to cause long-lasting and large-scale changes in retinal processing and is central to the process of light adaptation (Roy & Field, 2019; Witkovsky, 2004). For example, dopamine mediates the light-evoked uncoupling of AII-AC gap junction networks via activation of D1Rs (Bloomfield & Völgyi, 2009; Hampson, Vaney, & Weiler, 1992; Kothmann, Massey, & O’Brien, 2009) which is critical to increasing the signal-to-noise ratio of the ON pathway and allows for improved visual acuity and contrast perception at increased ambient luminance, especially during transitions from scotopic to photopic conditions (Jackson et al., 2012; Roy & Field, 2019). The knowledge of how dopamine modulates crossover inhibition via glycine release, however, was limited to early suggestive studies (Jensen, 1992; Jensen & Daw, 1986; Pycock & Smith, 1983), but progress has been made more recently. In a study by Mazade et. al (2019), D1 receptor activation was shown to reduce light-evoked and spontaneous inhibition of OFF-CBCs to light-adapted levels (Mazade, Flood, & Eggers, 2019). Although the authors pointed to reduced glycine release by AII-ACs as the most likely causal explanation for their results, this hypothesis, as well as the underlying presynaptic mechanism, remained to be tested.

Since dopamine can change the physiology of all major cell types in the retina, including the DA-ACs themselves via D2 autoreceptors (Nguyen-Legros, Versaux-Botteri, & Vernier, 1999; Margaret L. Veruki, 1997), the interpretation of dopamine effects is challenging at both single cell and system levels. Gap junction networks between different cell types further compound the problem. Here, we use two different strategies to help unravel dopamine’s role in AII-AC signaling. First, we examined the effects of exogenous dopamine and D1R agonists/antagonists on presynaptic AII-AC membrane voltage and spiking properties in conventional Ames medium. Second, we adopted a reduced circuit strategy using synaptic blockers and the mGluR6 agonist L-AP4 to isolate the effects of D1 modulation on AII-AC intrinsic properties and spontaneous glycinergic IPSCs in type-2 OFF-CBCs, in both wild type (*wt*) and Cx36 knockout (*Cx36^−/−^*) mice. We found that dopamine and a D1R agonist both hyperpolarize about half of AII-ACs, but on a different time-scale and with different effects on spiking. In presynaptic AII-ACs, blockade of D1Rs caused membrane voltage depolarization and a reduction in both the R_input_ and intrinsic excitability, while in postsynaptic type-2 OFF-CBCs, we observed an increase in glycinergic spontaneous IPSCs. These effects were absent in *Cx36^−/−^* animals and we provide an argument that they are primarily mediated by gap junctions between AII-ACs and ON-CBCs.

## Materials and Methods

### Rodents and retinal slices

Animal procedures were approved by the Institutional Animal Care and Use Committee (IACUC) of the Oregon Health and Science University, in agreement with National Institutes of Health (NIH) guidelines and polices. Experiments were performed using C57BL/6 background mice (The Jackson Laboratory) of either sex, postnatal day 28-60. For recordings in OFF-CBCs, transgenic Synaptogamin2-EGFP mice (*Syt2-GFP*; MMRRC) were used, which specifically labels type-2 OFF-CBCs (which receive the majority of the glycinergic inputs from AII-ACs (Graydon et al., 2018; Meadows, Balakrishnan, Wang, & von Gersdorff, 2021). For convenience, we refer to “type-2 OFF-CBCs” simply as “OFF-CBCs” in the text. Connexin-36 knockout mice (*Cx36^−/−^*; generated as described by (Hormuzdi et al., 2001) were used in some experiments with AII-ACs (where indicated), and the two strains were bred to obtain *Syt2-Cx36^−/−^* animals.

Animals were deeply anesthetized using isoflurane (Novaplus) and the eyes removed after death by cervical dislocation. Dissection to extract the retina was performed in cold (∼4° C), carbogenated (95% O_2_, 5% CO_2_) Ames medium (US Biologicals) or cutting solution (described next). Retinas were then embedded in low melting point agarose (σ Type VIIA, 3% in Ames medium) and 250 µm slices were obtained using a vibratome (Leica Microsystems, VT1200S) in cold (∼4° C), carbogenated low-Na^+^ sucrose-based cutting solution composed of (in mM): 210 sucrose, 35 NaHCO_3_, 20 HEPES, 10 glucose, 6 MgCl_2_, 3 Na-pyruvate, 2.5 KCl, 1.25 NaH_2_PO_4_, and 0.5 CaCl_2_, buffered to pH 7.4 with NaOH. Retinal slices were then stored at room temperature in continuously carbogen-bubbled Ames solution for later transfer to the recording chamber. A peristaltic pump was used to continuously perfuse (at ∼2mL/min) carbogen-bubbled Ames solution (containing 1.15 mM Ca^2+^; (Ames & Nesbett, 1981) over the slices in the recording chamber. Our working volume of external (Ames) solution was 15 - 50 mL (recirculated by the pump until the end of each experiment). Recordings were performed at near physiological temperature (32-34°C) by use of an in-line heater (Warner Instruments), under dim red room light, low photopic (mesopic) conditions.

### Cell visualization and microscopy

Retinal slices were visualized after recording with differential interference contrast (DIC) microscopy and post-hoc epifluorescence microscopy using a water immersion objective lens (60X, Olympus). For some recordings, we used a fixed-stage, microscope Carl Zeiss Axioskop 2FS microscope connected to a Retiga SRV digital camera (Q-imaging). For the remaining experiments, we used an upright Olympus BX51WI microscope with a motorized stage connected to a Hamamatsu ORCA digital camera (Hamamatsu Photonics). AII-ACs were identified by their morphology, sublaminar position in the inner nuclear layer (INL) and electrophysiological properties (e.g., spikelet firing). Alexa 594 dye (10µM, Invitrogen) was used to capture epifluorescence images of the cell’s morphology before or after recording using either QCapture (Q-Imaging) or cellSense (Olympus) imaging software.

### Patch-clamp electrophysiology

Patch-clamp recordings were performed using double EPC-10/2 amplifiers controlled by the Patchmaster (2×91 version) software (HEKA Elektronik). Data were acquired at 20-100 KHz and low-pass filtered at 2.9 KHz. Borosilicate capillary glass (1B150F-4, World Precision Instruments) pipettes were pulled to 8-10 MΩ using a Narishige puller PP-830 (Tokyo).

### Recordings of AII-AC spiking

Whole-cell current-clamp recordings of AII-ACs spiking were performed in Ames solution (280 mOsm) using the following K-gluconate-based internal solution (in mM): 104 K-gluconate, 10KCl, 10 HEPES, 10 Na_2_-phosphocreatine, 4 ATP-Mg, 2 EGTA, 0.4 GTP-Na_2_, adjusted to a pH of 7.4 with KOH (275 mOsm). When explicitly noted, the AMPA/kainate receptor blocker CNQX (10µM), the NMDA receptor blocker DL-AP5 (50µM), the GABA_(A)_ receptor blocker gabazine (SR95531, 8µM) and the group-III metabotropic glutamate receptor (mGluR) agonist L-AP4 (10µM) were added to the external (Ames) solution to specifically isolate the AII-AC intrinsic properties, the electrical synapses between AII-ACs and ON-CBCs, the glycinergic synapses between AII-ACs and OFF-CBCs, and control for confounding variables that would prevent clear interpretations of dopamine’s effects. AII-ACs were set to a V_m_ of ∼−55 mV to prevent rundown of spikelet frequency and amplitude seen at more depolarized membrane potentials, which can most likely be attributed to Na^+^ channel inactivation. During recording, the V_m_ was allowed to fluctuate naturally. Previous studies in either light or dark-adapted retina reported an AII-AC resting V_m_ between −50 mV and −40mV (Bloomfield & Xin, 2000; Dunn, Doan, Sampath, & Rieke, 2006; Tian et al., 2010). Additionally, we noted that a V_m_ ∼ −55 mV is ideally suited for analysis of burst firing in AII-ACs. Experimental drugs were directly applied to the bath and continuously washed for at least 5 minutes. Most attempts of drug washout were unreliable and excluded from analysis because of the long time required due to the high affinity of the compounds. For some experiments, spikelets were recorded at ∼−50 mV, −55 mV and −60 mV, for 60 to 90 seconds each, before and during drug wash, to extract spiking properties as a function of V_m_ and detect changes in intrinsic excitability. V_m_ was measured as the peak (statistical mode) of the histogram (bin = 1 mV) of the membrane potential. Spikelets and bursts of AII-ACs were detected and analyzed offline using custom routines in IgorPro (Wavemetrics) software and Python scripts based on scipy.signal methods (code can be found at GitHub: https://github.com/ADagostin/Igor-Procedures).

The burst detection method was primarily based in abrupt changes in instantaneous spikelet frequency (Δf_insta_) combined with an intra-burst spikelet frequency threshold (f_threshold_). To illustrate, setting Δf_insta_ = 3 and f_threshold_ = 100 Hz means that a *potential* burst would be detected as an >3-fold change in f_insta_ and confirmation would follow only if f_threshold_ > 100 Hz. Values for Δf_insta_ ranged from 2 (cells with a mean frequency ≥ 120 Hz) to 4 (cells with a mean frequency ≤ 20 Hz) and f_threshold_ from 80 to 250 Hz. For each individual cell, the value of f_threshold_ corresponded to the second well defined peak on a f_insta_ (1/inter-event interval) histogram (bin = 3 Hz), which was lacking in cells with no burst.

### Current step recordings in AII-Acs

Whole-cell current-clamp recordings at different current injections were performed in Ames (280 mOsm) using the previously described K-gluconate-based internal solution (275 mOsm). AII-ACs were set to a V_m_ of ∼−55 mV and current was injected for one second from −200 pA to +100 pA in 50 pA intervals. Next, the synaptic blocker cocktail containing CNQX (10µM), DL-AP5 (50µM), gabazine (8µM) and L-AP4 (10µM) was continuously applied for 5 minutes and the current injection protocol was repeated. Resting V_m_ was calculated at 0 pA as the statistical mode of a V_m_ histogram (bin = 1 mV) and input resistance was calculated as the slope (linear regression) of the cell’s voltage-current relationship.

### Postsynaptic recordings of sIPSCs from OFF-CBC

Whole-cell voltage-clamp recordings of type-2 OFF-CBCs spontaneous IPSCs (sIPSCs, holding = 0 mV) from *Syt2-EGFP* mice were performed in external Ames solution (280 mOsm) using the following Cs-based internal solution (in mM): 116 Cs-gluconate, 10 HEPES, 5 TEA-Cl, 5 Na_2_-phosphocreatine, 4 ATP-Mg, 4 EGTA, 4 QX314 Chloride, 0.5 GTP-Na_2_, adjusted to a pH=7.4 with CsOH and osmolarity of 275 mOsm. Glycinergic sIPSCs were isolated by use of the synaptic blockers CNQX (10µM), DL-AP5 (50 µM) and SR95531 (8 µM) in the external (Ames) solution. In addition to controlling for variations in ambient light, L-AP4 was also used to reduce very high frequency, extensively overlapping sIPSCs. Experimental drugs were directly applied to the bath and continuously washed for at least 5 minutes. Strychnine (2 µM) was applied at the end of some experiments to confirm that all sIPSCs were glycinergic and to obtain noise levels. The glycinergic sIPSCs were detected and analyzed offline using custom routines in IgorPro (Wavemetrics) software and Python scripts based on scipy.signal methods (code can be found at GitHub: https://github.com/ADagostin/Igor-Procedures). Overlapping sIPSCs were particularly challenging to detect with sliding template (Clements & Bekkers, 1997) methods, which often missed around 20% of the events, including high amplitude ones. As an alternative, a method based on peak (local maximum) detection algorithms was used for that purpose, but not for amplitude or kinetics analysis. sIPSCs amplitude distributions were generated using a 3 pA bin, and inter-event distributions using a 10 ms bin. Data fittings were all performed in Prism 9 (GraphPad).

### Pharmacology

Drugs and salts stock solutions were prepared with ultra-pure water (Mili-Q, Millipore) and stored in freezer at −20° C. Synaptic blockers CNQX, DL-AP5, gabazine (SR95531) and L-AP4, were all purchased from Tocris Bioscience. TTX and strychnine were obtained from Sigma Aldrich. Bath-applied drugs used to investigate dopaminergic modulation were dopamine hydrochloride (Sigma), D1R agonist SKF38393, and D1R antagonists SCH23390 and SKF83566 (all from Tocris). During experiments, dopamine stock aliquots and the syringes containing internal solution were kept in an ice box covered with aluminum foil to avoid degradation by temperature or light. Ultra-pure salts for internal and external solutions were all purchased from Sigma.

### Experimental design and statistical analysis

Shapiro-Wilk tests were used to verify normality of datasets from all experiments. Two-tailed paired *t*-tests were applied to compare pairwise data. Repeated measures one-way ANOVA were performed to compare paired groups followed by *post-hoc* Sídák’s multiple comparisons tests. All statistical tests were performed in Prism 10 (GraphPad) software. Data and error bars are reported as mean ± standard deviation. When drugs were washed, only one cell per slice was recorded or used for analysis. We adopted *α* = 0.05 for all tests and statistical significance is denoted by asterisks as follows: *: *p* < 0.05; **: *p* < 0.01; ***: *p* < 0.001; ****: *p* < 0.0001. Statistically insignificant results are labeled by “ns”.

## Results

### Spiking properties of AII-ACs depend strongly on membrane voltage

We found that the spiking properties of AII-ACs were strongly dependent on membrane voltage (V_m_). In current-clamp mode, we recorded the spikelets at different membrane voltages from −65 mV to −45 mV. At more hyperpolarized potentials, such as −65 to −60 mV, AII-AC spiking was characterized by low frequency, high amplitude bursts which could be interspersed with occasional, large amplitude single spikelets (**Figure 1 A*(i)***). The underlying burst’s waves could be long enough to contain several superimposed spikelets or could be very short and just contain doublets or triplets. At −55 mV, overall spikelet frequency increased with a predominance of single spikelets (**Figure 1 A*(ii)***), whereas at more depolarized potentials, such as −45 mV, spiking was characterized by high-frequency, low-amplitude singlet trains without bursts (**Figure 1 A*(iii)***). The relationship between AII-AC spiking behavior and membrane voltage were quantified in **Figure 1 B**. Spikelet frequency at −60 mV was 38.3 ± 16.4 Hz and increased approximately linearly to 118.2 ± 20.9 Hz as the membrane depolarized to −45 mV (**Figure 1 B*(i)***). These values reflected overall spikelet frequency which included spikelets within bursts. Burst frequency, on the other hand, decreased as membrane voltage depolarized, starting at 7.8 ± 3.9 Hz at −60 mV and decreasing to around 1.3 ± 0.9 Hz by −45 mV (**Figure 1 B*(ii)***). The burst frequency standard deviation was highest at −55 mV, when diverse spiking patterns were more readily recognizable. Spike amplitude decreased from 11.8 ± 3.0 mV to 8.6 ± 2.4 mV as the AII-ACs depolarized (**Figure 1 B*(iii)***). This difference would be greater if we had considered only single spikelets, as bursting spikelets typically have smaller amplitudes. We also obtained a voltage-current relationship (VI curve) for each cell from which we calculated the input resistance (R_input_) at both −55 mV and resting membrane potential (V_rest_). The R_input_ at V_rest_ (−32.5 ± 7.2 mV) was lower than at −55 mV (312 ± 76 MW vs 199 ± 52 MW; *p* = 0.0001; **Figure 1B*(v)***), a result that was clear when examining the mean VI curve shown in **Figure 1B*(iv)***. This data suggests that AII-ACs have a very depolarized V_m_ with low R_input_ under normal Ames media and dim photopic conditions, so tonic glycinergic output to downstream OFF-CBCs may be very high.

**Figure 1.**
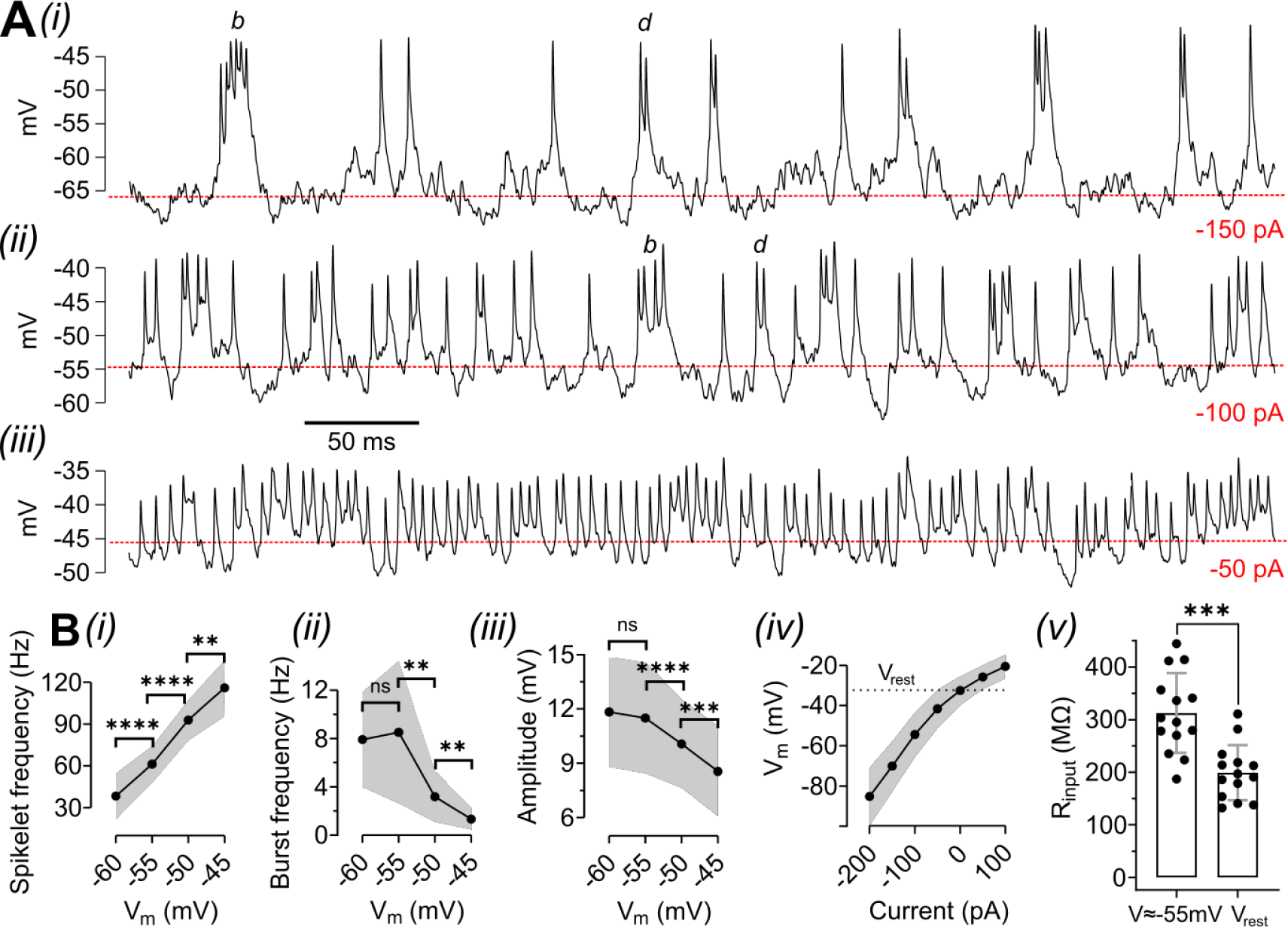
Electrophysiological properties of AII-ACs. **(A)** Current-clamp recordings of spiking behavior of AII-ACs at different Vm. ***(i)*** At hyperpolarized potentials (around −65 to −60 mV) AII-AC’s typically exhibited prominent bursting behavior, with some cells also presenting large amplitude single spikelets. ***(ii)*** Single spikelets were more frequent than bursts at around −55 mV, when a greater diversity of spiking patterns was observed. Letters “*b*” and “*d*” indicate examples of bursts and doublets. ***(iii)*** Spiking at more depolarized potentials (around −45mV) was characterized by high frequency, small amplitude single spikelets. Red dotted lines correspond to the Vm mode (peak value of a Vm histogram). **(B)** Spikelet frequency ***(i)***, burst frequency ***(ii)*** and spikelet amplitude ***(iii)*** as a function of Vm. ***(iv)*** The current-voltage relationship of AII-ACs. The dotted line shows the resting Vm (Vrest), when no current was injected (0 pA). ***(v)*** Like other parameters measured, the variance of AII-AC Rinput was high, both at −55 mV and Vrest. Rinput was significantly lower when measured at Vrest. *n* = 14 cells. Shaded areas and error bars correspond to the mean ± standard deviation. Statistical significance was determined using paired *t*-tests or repeated measures one-way ANOVA followed by Sídák’s multiple comparisons; ns: *p* > 0.05; **: *p* < 0.01; ***: *p* < 0.001; ****: *p* < 0.0001.

### Dopamine and a D1R agonists hyperpolarize a subset of AII-ACs

There are many reports of how dopamine modulates the excitability of RGCs (Chen & Yang, 2007; Cui et al., 2017; Hayashida et al., 2009; Ogata, Stradleigh, Partida, & Ishida, 2012), but no studies have yet demonstrated how dopamine changes AII-AC membrane potential (V_m_) and spikelet firing properties. Here we investigated the effects of exogenous dopamine and a D1R agonist on AII-AC membrane potential and spiking properties. Current was injected to keep cells at approximately −55 mV, then membrane voltage and spikelets were recorded before and during bath application of 10 mM dopamine or the D1R agonist SKF38393. We found that exogenous dopamine resulted in relatively fast membrane voltage hyperpolarization and a noticeable reduction in spikelet frequency (**Figure 2 A*(i)***), while SKF38393 resulted in a slower hyperpolarization without the corresponding change in spikelet frequency (**Figure 2 A*(ii)***). Interestingly, only a subset of AII-ACs responded to exogenous dopamine or the D1R agonist, with the proportions being 53% (8 out of 15 cells) and 44% (7 out of 16 cells) respectively (**Figure 2 B**).

**Figure 2.**
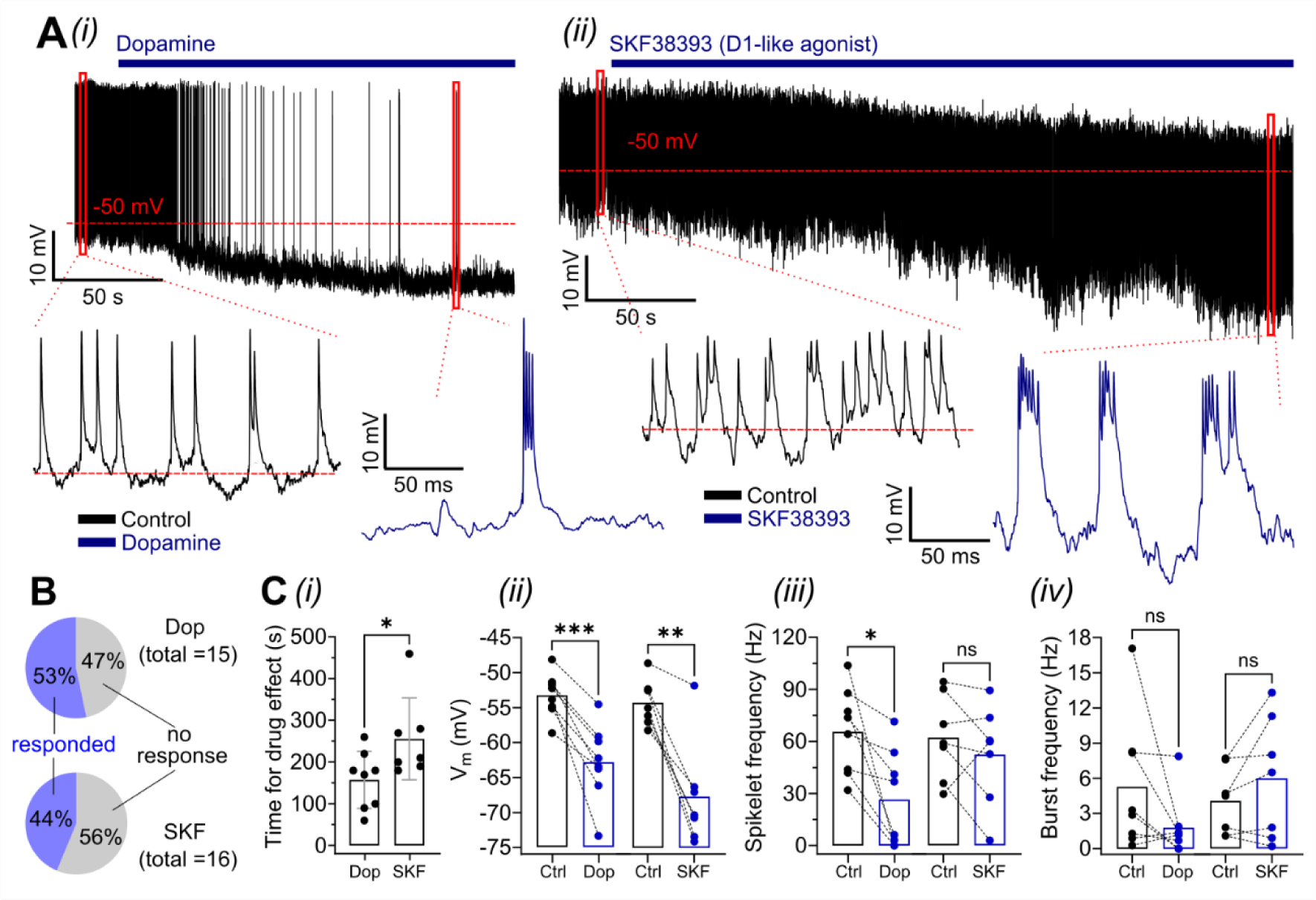
Dopamine and a D1R agonist both hyperpolarized AII-ACs. **(A)** Example current-clamp recordings from AII-ACs showing the hyperpolarization caused by bath application of 10 μM dopamine ***(i)*** and the D1R agonist SKF38393 ***(ii)***. Note that dopamine, but not the D1R agonist, strongly reduced spiking, even though the hyperpolarization was less pronounced. Note also that the response to the D1R agonist was slower. *Insets* show AII-ACs spiking in control (*black*) and in treatment (*blue*) conditions. **(B)** About half of the cells recorded did not respond to either drug (supplemental material). **(C)** Comparison of dopamine and D1R agonist effects. ***(i)*** Time for full effect (no further change in parameters) to be observed. AII-ACs membrane potential ***(ii)***, spikelet frequency ***(iii)*** and burst frequency ***(iv)*** in control (*black*) and after drug wash (*blue*). Dopamine: *n* = 8 cells; SKF38393: *n* = 7 cells. Circles in bar graphs represent single cells and error bars represent the mean ± standard deviation. Statistical significance was determined using paired *t*-tests; ns: *p* > 0.05; *: *p* < 0.05; **: *p* < 0.01; ***: *p* < 0.001.

**Figure 2 C** quantifies the effects of dopamine and SKF38393 on membrane potential and spiking properties. The estimated time for the full effect (no further change in V_m_) was 157 ± 69 s for dopamine and 256 ± 98 s for SKF38393 (*p* = 0.0407; **Figure 2 C*(i)***). Exogenous dopamine hyperpolarized AII-AC membranes from a baseline V_m_ of −53.1 ± 3.1 mV to −62.9 ± 5.5 mV (*p* = 0.0007), with the strongest responding cell hyperpolarizing 19.5 mV. For SKF38393 we observed a change in V_m_ from a baseline of −54.2 ± 3.3 mV to −67.8 ± 7.6 mV (*p* = 0.0011), with the strongest responding cell hyperpolarizing 20.9 mV. Mean spikelet frequency dropped with exogenous dopamine from 65.7 ± 24.9 Hz to 26.6 ± 27.9 Hz (*p* = 0.0124), but did not change with the application of SKF38393: 62.3 ± 24.8 Hz to 52.4 ± 28.9 Hz (*p* = 0.2957). In 2 out of 8 cells, dopamine completely and permanently inhibited firing even before hyperpolarization peaked. We did observe a clear drop in spikelet frequency in 3 out of 7 cells following application of SKF38393, but this effect was accompanied by a very strong hyperpolarization. Interestingly, no cell stopped firing and, on the contrary, bursting activity was still present at a V_m_ near or below −70 mV, a behavior never observed in control conditions in any experiment. Burst frequency was not changed by either dopamine or SKF38393 (dopamine: 5.3 ± 5.7 Hz to 1.8 ± 2.6 Hz, *p* = 0.1106; SKF38393: 4.1 ± 2.9 Hz to 6.0 ± 5.2 Hz, *p* = 0.1523; **Figure 2 C*(iv)***), although dopamine dramatically reduced this bursting in some cells. Overall, these results suggest that dopamine reduce AII-AC excitability through a combination of local and circuit-level mechanisms, potentially involving synaptic release changes and activation of D1R and D2R in other cells, while specific activation of D1-like receptors by SKF38393 causes an apparent increase in AII-AC excitability, because it hyperpolarizes the AII-AC but does not reduce spike rates and spiking can be seen at V_m_ well below the subthreshold level in control conditions.

Importantly, we also performed statistical analysis on all cells grouped together without splitting the dataset into responding and non-responding groups. Hyperpolarization of AII-ACs by both dopamine and SKF38393 can still be observed (dopamine: *p* = 0.0067; SKF38393: *p* = 0.0228). However, except for a reduction in spikelet frequency by dopamine (*p* = 0.0099), we found no other statistically significant effects on other spiking properties by either drug. Additionally, we found no differences in the parameters measured when comparing responding and non-responding groups in control conditions. A cell was considered responsive if a 2σ change in at least two parameters was observed, with σ being the standard deviation in control conditions calculated for all cells.

### A D1R antagonist depolarized a subset of AII-ACs

Since both exogenous dopamine and the D1R agonist SKF38393 were without effect in approximately half of the cells recorded, we considered the hypotheses that D1Rs are either tonically activated or saturated by endogenous dopamine, or that a fraction of AII-ACs do not express functional D1Rs. To discriminate between them, we decided to test the effects of the D1R antagonist SKF83566 on AII-ACs. Just like in the previous experiment, current was injected to keep cells at approximately −55 mV, then membrane voltage (V_m_) and spiking properties were recorded before and during a bath application of 5 mM SKF83566. In favor of the saturation hypothesis, we observed a slow (271 ± 20 s) and modest depolarization in only some AII-ACs (**Figure 3 A*(i)***), while in others, the V_m_ was unchanged (**Figure 3 A*(ii)***). In this experiment, exactly 50% of AII-ACs responded to treatment (7 out of 14 cells; **Figure 3 B**). In the responding group, SKF83566 depolarized the cells from −54.6 ± 2.3 mV to −50.0 ± 2.6 mV (*p* = 0.0005), while in the non-responding group, V_m_ was −54.4 ± 4.9 mV in control and −54.6 ± 4.7 mV after treatment (*p* = 0.7235; **Figure 3 C*(i)***). SKF83566 had no effect on mean spikelet frequency in cells classified as either responding or non-responding (responding: 73.1 ± 15.8 Hz to 64.5 ± 14.5 Hz, *p* = 0.2542; non-responding: 68.1 ± 29.4 Hz to 68.4 ± 33.9 Hz, *p* = 0.9650; **Figure 3 C*(ii)***). This suggests that D1R blockade decreases the excitability of AII-ACs, otherwise the depolarization caused by the D1R antagonist should be accompanied by an increase in spikelet frequency (see **Figure 1 B*(i)***). This is consistent with the previous experiment, where the D1R agonist increased the excitability of AII-ACs. SKF38566 also had no effect on mean burst frequency (responding: 7.6 ± 6.2 Hz to 4.6 ± 3.0 Hz, *p* = 0.0.0778; non-responding: 3.8 ± 3.9 Hz to 2.7 ± 2.7 Hz, *p* = 0.1723; **Figure 3 C*(iii)***), though 6 out of 7 cells in the responding group did show a reduction after drug application, with two of them showing more than a 50% drop in the mean value. Finally, a small decrease in the spikelet amplitude of the responding cells was observed (8.9 ± 3.8 mV to 7.9 ± 4.1 mV, *p* = 0.0479), which was expected as a consequence of the depolarization.

**Figure 3.**
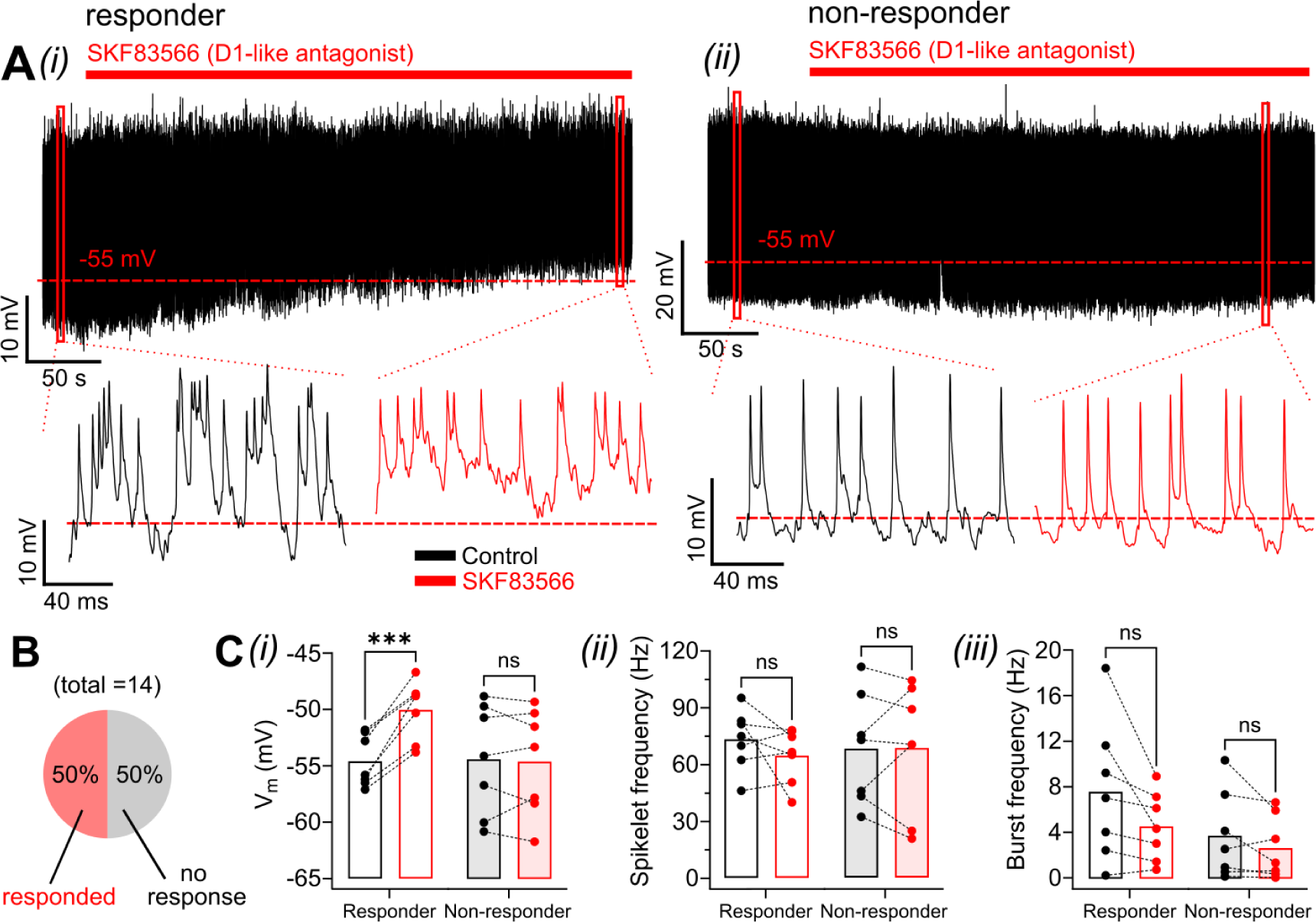
A D1R antagonist depolarized about half of AII-ACs, suggesting tonic activation of D1Rs by endogenous dopamine in a subset of AII-Acs. **(A)*(i)*** Current-clamp recording from an AII-AC showing the slow depolarization caused by bath application of the D1R antagonist SKF83566 (5 μM). ***(ii)*** An AII-AC that did not respond to SKF83566. *Insets* show AII-ACs spiking in control (*black*) and in treatment (*red*) conditions. Note that although the cell depolarizes in *(i)*, the spiking slightly decreases, which suggests decreased excitability. **(B)** Fraction of cells that responded and did not. AII-ACs membrane potential ***(i)***, spikelet frequency ***(ii)*** and burst frequency ***(iii)***, measured in control (*black*) and after drug wash (*red*). *n* = 7 cells for both groups. Circles in bar graphs represent single cells. Statistical significance was determined using paired *t*-tests; ns: *p* > 0.05; ***: *p* < 0.001.

We also observed SKF83566 effects when all cells, both responding and non-responding, were grouped together. In addition to the V_m_ depolarization (*p* = 0.0180), a decrease in spikelet amplitude (*p* = 0.0224) and burst frequency (*p* = 0.0247) was found. Again, we found no differences in the parameters when comparing responding and non-responding groups in control conditions.

### A D1R agonist had no effect on AII-AC membrane potential or OFF-CBC sIPSCs in the presence of synaptic blockers

Next, we wanted to examine the consequences of D1R modulation of AII-AC excitability and membrane potential by examining AII output to downstream OFF-CBCs. Because all major cell types in the retina express dopamine receptors, and since our previous results could be influenced by light responses from photoreceptors or by glutamatergic and GABAergic synapses that also contain D1Rs, we designed an experimental condition to isolate, as much as possible, both the intrinsic properties of AII-ACs and glycinergic synapses to downstream OFF-CBCs (a reduced-circuit approach). To this end, the following experiments with the D1R agonist SKF38393 was performed in Ames solution containing the AMPA/kainate receptor blocker CNQX (10 µM), the NMDA receptor blocker DL-AP5 (50 µM), the GABA_(A)_ receptor blocker gabazine (8 µM), and the group-III mGluR agonist L-AP4 (10 µM). Furthermore, we also chose to perform non-paired recordings of postsynaptic OFF-CBCs to keep the intracellular composition of the presynaptic AII-ACs intact. While AII-AC lobular appendages form synapses with multiple OFF-CBC subtypes, synaptic targeting is heavily biased toward type-2 OFF-CBCs (also called CBC2s) (Graydon et al., 2018). We specifically targeted fluorescently labeled type-2 OFF-CBCs in mice expressing GFP under control of the synaptotagmin-2 (*Syt2*) promoter (*Syt-GFP*). By holding GFP-positive type-2s at 0 mV, we could record their positive spontaneous inhibitory post-synaptic currents (sIPSCs) as a proxy for AII-AC glycine release.

In our previous experiments, we could not distinguish between local (cell-specific) and circuit level effects of the D1R agonist on AII-ACs. By using the synaptic blocker cocktail, we expected to observe only the former, so that glycinergic output of AII-ACs could be more directly correlated to changes in their intrinsic properties caused exclusively by activation of their own D1Rs. To our surprise, SKF38393 no longer affected AII-AC membrane voltage or spiking properties (**Figure 4A**), and, accordingly, also had no effect on downstream OFF-CBC glycinergic sIPSCs (**Figure 4B**). AII-AC V_m_ (−52.7 ± 1.4 mV to −52.8 ± 1.2 mV, *p* = 0.8519; **Figure 4C*(i)***), spikelet frequency (62.9 ± 34.2 Hz to 58.4 ± 31.5 Hz, *p* = 0.2360; **Figure 4C*(ii)***), burst frequency (2.3 ± 3.0 Hz to 2.4 ± 3.7 Hz, *p* = 0.9134; **Figure 4C*(iii)***), and spikelet amplitude (6.9 ± 1.7 mV to 6.9 ± 1.5 mV, *p* = 0.8098; **Figure 4D*(iv)***) remained unchanged after drug wash. The same applied to spike and burst frequencies analyzed as a function of V_m_ (2-way ANOVA; *p* = 0.0573 and *p* = 0.6787, respectively), meaning AII-AC intrinsic excitability was unaffected. R_input_ (362 ± 128 MW to 345 ± 132 MW, *p* = 0.2283; **Figure 4D*(i)***) and the AII-AC IV curve (2-way ANOVA; *p* = 0.5525; **Figure 4D*(ii)***) also remained the same after SKF38393 application.

**Figure 4.**
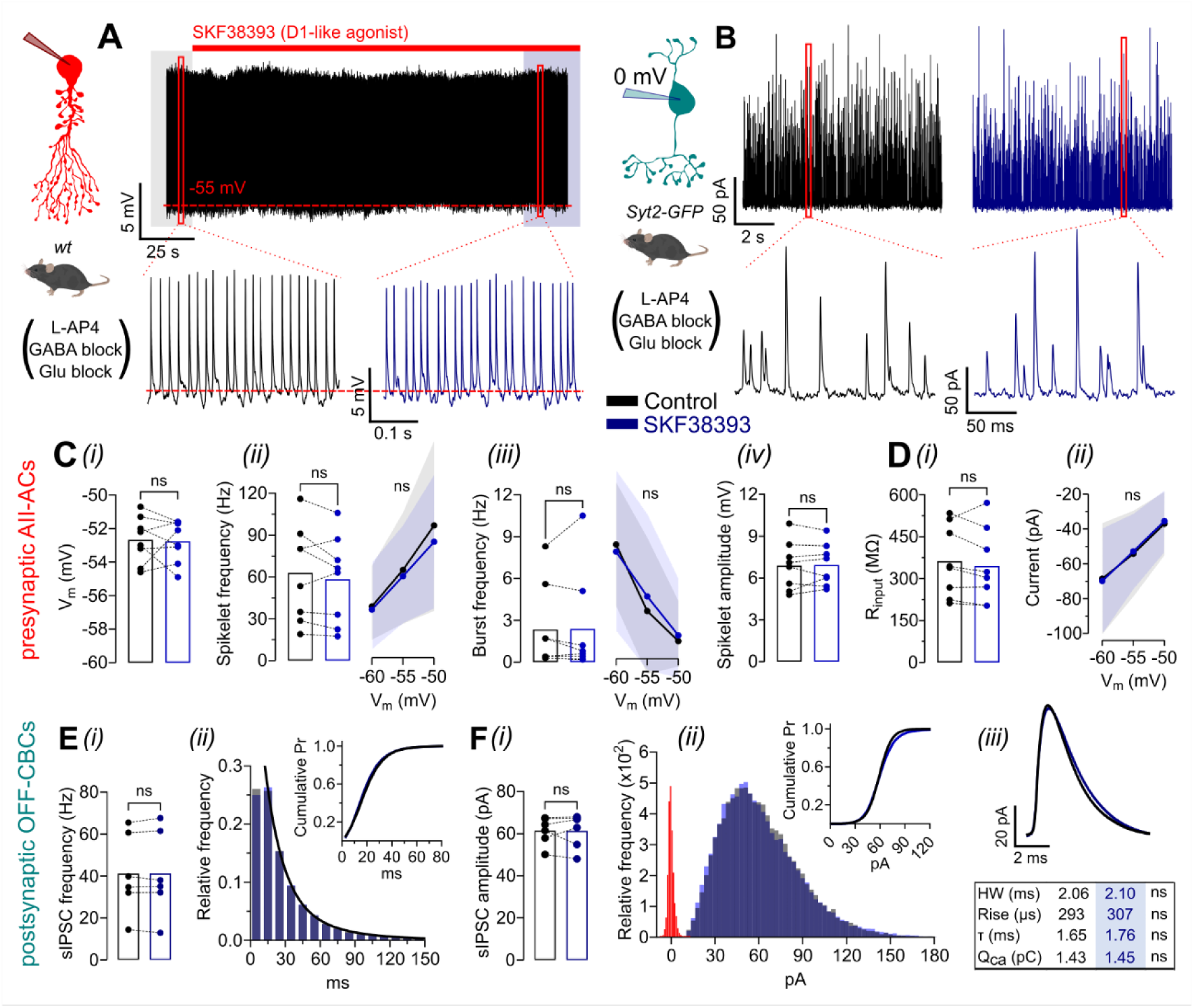
A D1R agonist had no effect on AII-AC membrane potential and OFF-CBC sIPSCs in the presence of synaptic blockers. **(A)** Example current-clamp recording of an AII-AC showing that bath application of the D1R agonist SKF38393 (10µM) does not change its Vm or spiking (*inset*). **(B)** Example voltage-clamp recording (holding = 0 mV) of an OFF-CBC showing that the agonist also does not affect glycinergic spontaneous IPSCs (sIPSCs). **(C)** AII-AC’s membrane potential (Vm) ***(i)***, spikelet frequency ***(ii)***, burst frequency ***(iii)***, and spikelet amplitude ***(iv)*** measured in control and after SKF38393. Right sides of ***(ii)*** and ***(iii)*** show spikelet and burst frequencies as a function of Vm. **(D)** Rinput *(**i)*** and IV curve ***(ii)*** of AII-ACs in control and after the D1R agonist. Note the absence of any significant effect on all parameters measured. **(E)** OFF-CBC sIPSC mean frequency ***(i)*** and sIPSC inter-event interval (IEI) distribution ***(ii)*** with corresponding cumulative probability plot (*inset*) in control and after SKF38393. The same double exponential decay fitted both IEI distributions. **(F)** sIPSC mean amplitude ***(i)*** and sIPSC amplitude distribution (***ii***; noise in red) with cumulative probability plot (*inset*) before and after SKF38393. ***(iii)*** Mean sIPSC waveforms for each experimental condition. Table compares the kinetics and charge of the mean sIPSC for each individual cell. AII-ACs: *n* = 6 cells; OFF-CBCs: *n* = 7 cells. Ctrl: 8410 ± 3700 events per cell, SKF38393: 8380 ± 4630 events per cell. Circles in bar graphs represent single cells. Statistical significance was determined using paired *t*-tests and repeated measures one-way ANOVA; ns: *p* > 0.05.

In the postsynaptic type-2 OFF-CBCs, SKF38393 had no effect on sIPSC frequency, either when analyzed by mean frequency (41.2 ± 19.1 Hz vs 41.2 ± 20.2 Hz, *p* = 0.9718; **Figure 4E*(i)***) or inter-event interval (IEI) (**Figure 4E*(ii)***). The IEI distributions for both control and SKF38393 conditions were fit with the same double exponential decay [t_fast_ = 15.2 ms (83%), t_slow_ = 41.7 ms, *p* = 0.9914, extra sum-of-squares *F*-test], and the cumulative probability curves (inset) were nearly identical. Additionally, we found no change in mean sIPSC amplitude (61.4 ± 6.6 pA to 61.3 ± 8.1 pA, *p* = 0.9654; **Figure 4F*(i)***). The amplitude distributions and cumulative probability curves were nearly overlapping before and after application of SKF38393 (**Figure 4F*(ii)***). We also compared the kinetics and charges of each cell’s mean sIPSC waveform before and after SKF38393 but found no difference (half-width: *p* = 0.5704; rise time: *p* = 0.1280; t_decay_: *p* = 0.0820; charge: *p* = 0.7759; mean values are reported in **Figure 4F*(iii)***). Overall, these results indicate that, in the presence of the synaptic blocker cocktail, D1Rs are most likely saturated so that no changes in AII-AC intrinsic properties or their spontaneous glycinergic output to OFF-CBCs could be detected following bath application of the D1R agonist.

### D1Rs modulate AII-AC membrane potential and excitability through gap junctions with ON-CBCs

In our experiments performed in dim photopic conditions and using Ames without the addition of any drugs, 44% of AII-ACs responded to the D1R agonist and 50% to the D1R antagonist, so it’s reasonable to supposed that endogenous dopamine levels are high enough to saturate or tonically activate D1Rs in about half of AII-ACs. In an unexpected way, supplementing the Ames solution with glutamatergic and GABAergic blockers, as well as L-AP4, obliterated any effect of the D1R agonist on either presynaptic AII-ACs or postsynaptic OFF-CBCs, suggesting that full saturation was reached. Independent of the mechanisms involved, if this hypothesis is correct, we should be able repeat the experiment from **Figure 3** and see effects of the D1R antagonist SKF83566 in a greater proportion of AII-ACs. As expected, SKF83566 depolarized AII-ACs (**Figure 5A*(i)***) in a very similar manner to that seen without the synaptic blocker cocktail (**Figure 3A*(i)***), except in this case, all cells responded to treatment.

**Figure 5.**
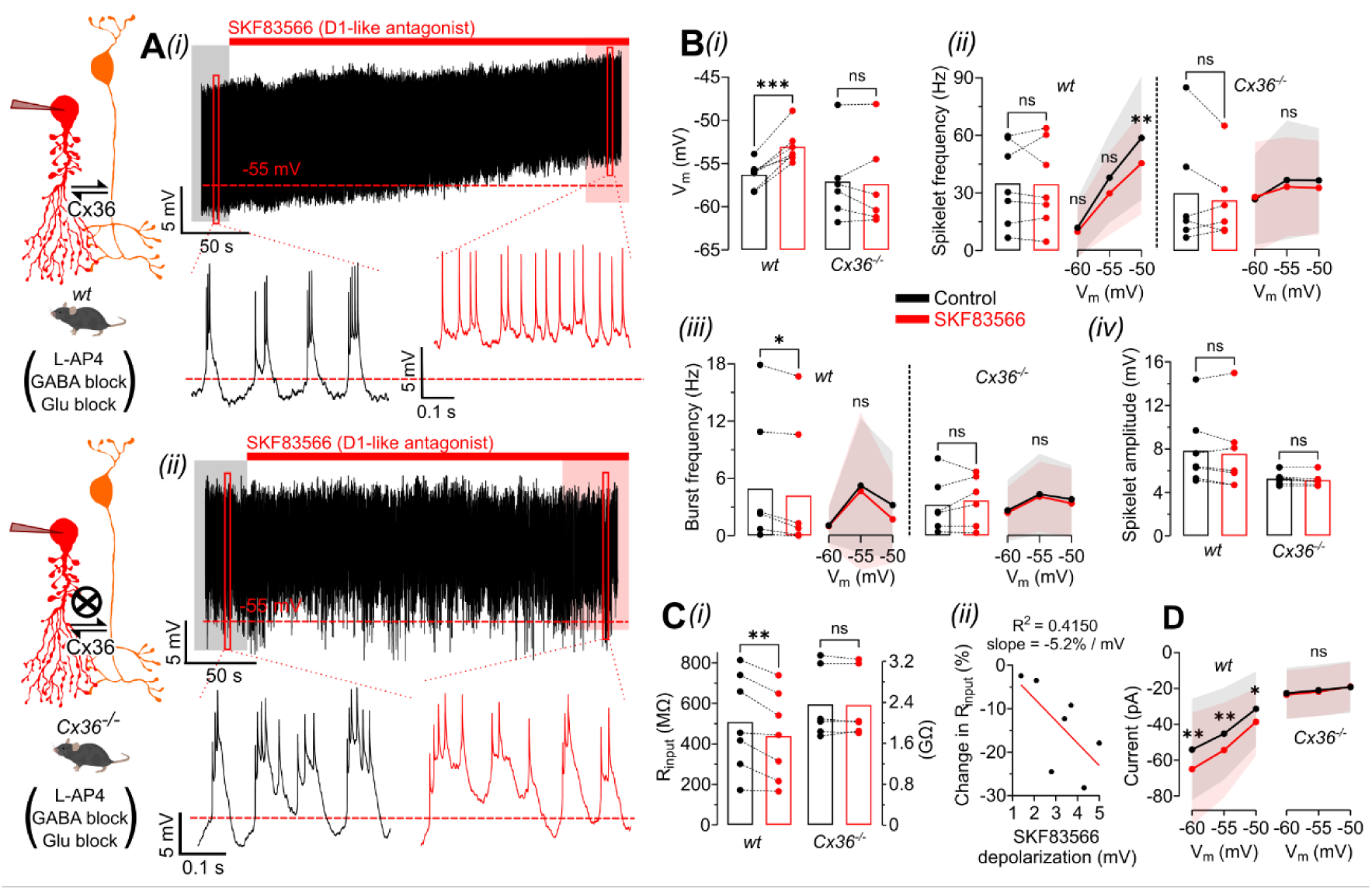
Blocking D1Rs depolarized AII-ACs in *wt* but not *Cx36^−/−^* animals. **(A)** Example current-clamp recordings contrasting how AII-ACs from *wt **(i)*** and connexin-36 knockout (*Cx36^−/−^*) ***(ii)*** mice respond to the D1R antagonist SKF83566 (5μM). In *wt* animals, bath application of the SKF83566 slowly depolarizes AII-ACs, while no clear effects were observed in cells of *Cx36^−/−^* animals. Experiments were performed in external solution containing a drug cocktail (round brackets) to isolate the AII-AC/ON-CBC system and control for variations in light conditions. *Insets* show AII-ACs spiking in control (*black*) and in treatment (*red*) conditions. **(B)** AII-ACs membrane potential ***(i)***, spikelet frequency ***(ii)***, burst frequency ***(iii)*** and spikelet amplitude ***(iv)***, measured in control (*black*) and after SKF83566 (*red*), in *wt* and *Cx36^−/−^* mice. Although relatively small, the depolarization caused by SKF83566 was consistent across all cells recorded in *wt* mice. Note that it was not accompanied by an increase in spikelet frequency, which would be expected based on the dependence of this parameter to the Vm of the cell. This later effect was a consequence of decreased *intrinsic* excitability, particularly around −50mV, as can be noticed on the spikelet frequency *vs* Vm curve showed on the right side of the bar graph. **(C)*(i)*** The input resistance (Rinput) decreased in cells from *wt* animals, but not in *Cx36^−/−^*. Note the units of MΩ for *wt* (left y-axis) and GΩ for *Cx36^−/−^* (right y-axis). This result is consistent with the idea that blockage of D1Rs increases gap-junction coupling between AII-ACs and ON-CBCs. ***(ii)*** A moderate correlation was found between the depolarization caused by SKF83566 and the percentage decrease in Rinput. **(D)** IV curve for control and treatment conditions. A downward shift was observed only in *wt* animals, suggesting transmission of positive/inward current from ON-CBCs to AII-ACs. *Wt*: *n* = 7 cells; *Cx36^−/−^*: *n* = 6 cells. Circles in bar graphs represent single cells. Shaded areas and error bars correspond to the mean ± the standard deviation. Statistical significance was determined using paired *t*-tests or repeated measures one-way ANOVA followed by Sídák’s multiple comparisons; ns: *p* > 0.05; **: *p* < 0.01; ***: *p* < 0.001.

It’s well-established that dopamine can modulate connexin-36 (Cx36) gap junctions between AII-ACs via D1 receptors (Bloomfield & Völgyi, 2009; Hampson et al., 1992), which activate G_αs_ proteins and trigger the canonical AC/cAMP/PKA signaling pathway. More specifically, there is evidence that D1R activation dephosphorylates Cx36 hemichannels in AII-ACs by protein phosphatase-2A (PP2A), causing the open probability of the channels to be reduced and therefore uncoupling the AII<-<AII networks (Kothmann et al., 2009; O’Brien, 2014). ON-BCs make gap junction connections with AII-ACs through both Cx36 and Cx45 and, relevant to our investigation, D1R antagonists have been shown to increase AII<->ON-CBC coupling (Urschel et al., 2006; Xia & Mills, 2004; Shubhash C. Yadav, Tetenborg, & Dedek, 2019). Therefore, the hyperpolarization of AII-ACs caused by exogenous dopamine and the D1R agonist SKF38393 (**Figure 2**) could be caused by uncoupling between AII-ACs and ON-CBCs, and the depolarization of AII-ACs caused by the D1R antagonist SKF83566 (**Figure 3 and Figure 5A**) could be the result of increased coupling.

To address this hypothesis, we also applied the D1R antagonist SKF83566 to Cx36 knockout (*Cx36^−/−^*) retinal slices in the presence of the synaptic blocker cocktail described previously (**Figure 5**) and compared them to our results from control (*wt*) animals. We found that the membrane depolarization seen in control AII-ACs was blocked in *Cx36^−/−^* mice (**Figure 5A*(ii)***). In control AII-ACs, V_m_ depolarized from −56.3 ± 1.5 mV to −53.1 ± 2 mV (*p* = 0.0005) whereas in *Cx36^−/−^* AII-ACs, V_m_ remained unchanged from −57.1 ± 4.7 mV to −57.4 ± 5.2 mV (*p* = 0.6510; **Figure 5B*(i)***). One cell out of the six recorded did depolarize, suggesting a secondary mechanism not involving Cx36 gap junctions. We also note that the amount of current injected to hold *Cx36^−/−^* AII-ACs at a V_m_ of ∼−55 mV was much smaller than in *wt* animals (*wt*: −57 ± 23 pA; *Cx36^−/−^*: −18 ± 7 pA; *p* = 0.0056). The spikelet frequency was unchanged in both *wt* and *Cx36^−/−^* AII-ACs (*wt*: 34.9 ± 21.3 Hz to 34.5 ± 22.3 Hz, *p* = 0.9006; *Cx36^−/−^*: 30.2 ± 30.0 Hz to 26.3 ± 20.8 Hz, *p* = 0.4196; **Figure 5B*(ii)***). However, when analyzed as a function of V_m_, spikelet frequency was reduced by SKF83566 at −50 mV, but only in *wt* AII-ACs (58.9 Hz to 45.7 Hz; *p* = 0.0038). This result directly supports our hypothesis from previous experiments performed in normal Ames (**Figures 2 and 3**) that D1R activation increases the excitability of AII-ACs and blockade of D1Rs reduces it. We note that, as synaptic blockers were used here, this conclusion refers only to the *intrinsic* excitability of AII-ACs. Average burst frequency was reduced in *wt* but not *Cx36^−/−^* AII-ACs (*wt*: 4.9 ± 6.8 Hz to 4.2 ± 6.7 Hz, *p* = 0.0175; *Cx36^−/−^*: 3.2 ± 2.9 Hz to 3.7 ± 2.6 Hz, *p* = 0.4191; **Figure 5B*(iii)***), an effect that is most likely a consequence of the depolarization itself, as no effect was observed when the parameter was compared as a function of V_m_ (2-way ANOVA; *wt*: *p* = 0.1130; *Cx36^−/−^*: *p* = 0.1407; **Figure 5B*(iv)***).

If the depolarization of AII-ACs is in fact caused by increased coupling between AII-ACs and ON-CBCs, as the V_m_ and spiking analysis between the two mouse strains strongly indicates, we would expect the input resistance (R_input_) measured in AII-ACs from *wt* animals to reduce after application of the D1R antagonist. We found that SKF83566 reduces R_input_ in *wt* but not *Cx36^−/−^* AII-ACs (*wt*: 508 ± 237 MW to 439 ± 218 MW, *p* = 0.0057; *Cx36^−/−^*: 2.4 ± 0.69 GW to 2.34 ± 0.67 GW, *p* = 0.7035; **Figure 5C*(i)***). Note that baseline R_input_ is significantly (about 4X) higher in *Cx36^−/−^* AII-ACs compared to *wt,* as expected in the absence of gap junctions. Moreover, when we plotted the percent change in R_input_ as a function of SKF83566 depolarization (**Figure 5C*(ii)***), we found a moderate negative correlation with an R^2^ value of 0.451. The significance of this finding is that it’s better explained by changes in gap junctions between AII-ACs and ON-CBCs than among AII-ACs. This is because coupled AII-ACs have been shown to display similar V_m_ and even synchronous oscillations (Hartveit & Veruki, 2012; Margaret Lin Veruki & Hartveit, 2002), so that changes in the conductance of their homotypic gap junctions would be expected to strongly affect only their R_input_. Also, supporting the ON-CBC coupling hypothesis, there was a downward shift in the IV curve in *wt* but not *Cx36^−/−^* AII-ACs (2-way ANOVA; *wt*: *p* < 0.0001; *Cx36^−/−^*: *p* = 0.6741; **Figure 5D**). This finding indicates the leakage of positive current through gap junctions from ON-CBCs to AII-ACs that is absent when coupling is knocked out. Taken together, these results provide evidence that D1R modulation of junctions with ON-CBCs plays an important role in setting the V_m_ and intrinsic excitability of AII-ACs.

### Blockade of D1Rs increases glycinergic transmission to OFF-CBCs in a gap junction dependent manner

To examine how D1R modulation of AII-AC V_m_ and excitability affects the glycinergic transmission to type-2 OFF-CBCs, we repeated the experiments in **Figure 4**, but now using the D1R antagonist SKF83566 instead of the D1R agonist (which showed no effect). We also developed the *Syt2-Cx36^−/−^* mouse strain to test for presynaptic mechanisms involving Cx36 gap junctions. If AII-ACs depolarize in response to D1R antagonists in a gap junction-dependent manner, an increase in glycinergic sIPSCs in the postsynaptic OFF-CBCs would be expected in the *Syt2-GFP* but not the *Syt2-Cx36^−/−^* animals. This is exactly what we found, as can be seen in voltage-clamp recordings in **Figure 6A*(i)*** and **Figure 6A*(ii)***. sIPSC frequency increased in *Syt2-GFP* from 33.3 ± 16.3 Hz to 44.3 ± 20.5 Hz (*p* = 0.0195) but was unchanged in *Syt2-Cx36^−/^* (11.2 ± 8.5 Hz to 11.4 ± 8.3 Hz, *p* = 0.8422; **Figure 6B*(ii)***). Interestingly, the baseline tonic inhibition from AII-ACs to OFF-CBCs was already strongly reduced in the *Syt2-Cx36^−/−^* OFF-CBCs (*p* = 0.0249, see **Figure 6A insets** for example traces), potentially due to the more hyperpolarized resting V_m_ in gap junction knockout AII-ACs. This scenario perfectly agrees with the conclusions from the previous experiment that increased coupling with ON-CBCs sets V_m_ in AII-ACs to more depolarized values. In **Figure 6C*(i)***, we show a time-series plot of cumulative sIPSCs for both strains, where a gradual increase in sIPSC frequency is seen in *Syt2-GFP* OFF-CBCs as a positive deflection beginning approximately 30 seconds after SKF83566 reached the bath. Interestingly, the increase in sIPSC frequency displayed a time-course very similar to that of the gradual depolarization of AII-ACs following SKF83566 application (**Figure 3A*(i)* and Figure 5A*(i)***). This drug-induced increase in sIPSC frequency in *Syt2-GFP* OFF-CBCs is also evident when we compare the inter-event interval (IEI) distributions in control vs treatment conditions: a double-exponential fit returns faster time constants after SKF83566 application [control: τ_fast_ = 22.1 ms (85%), τ_slow_: 49.7 ms; SKF83566: τ_fast_ = 10.6 ms (84%), τ_slow_: 35.6 ms; *p* < 0.0001, extra sum-of-squares *F*-test; left panel of **Figure 6C*(ii)***]. This difference can also be readily seen as a clear leftward shift in the corresponding cumulative probability plot (insert in left panel of **Figure 6C*(ii)***). In contrast, in *Syt2-Cx36^−/−^*animals, the IEI distributions for each experimental condition were both fitted by the same single-exponential and no shift in the cumulative probability plot is observed (τ = 61.5 ms, *p* = 0.9687, extra sum-of-squares *F*-test; right panel of **Figure 6C*(ii)***). Finally, sIPSC mean amplitude and waveform (not shown) did not change due to the application of SKF83566 in either *Syt2-GFP* (65.9 ± 9.6 pA to 64.7 ± 10.5 pA, *p* = 0.5109) or *Syt2-Cx36^−/−^* (48.1 ± 4.1 pA to 46.1 ± 4.2 pA, *p* = 0.3493; **Figure 6B*(i)***), suggesting that the drug is acting primarily on presynaptic AII-ACs and does not significantly affect postsynaptic mechanisms in OFF-CBCs.

**Figure 6.**
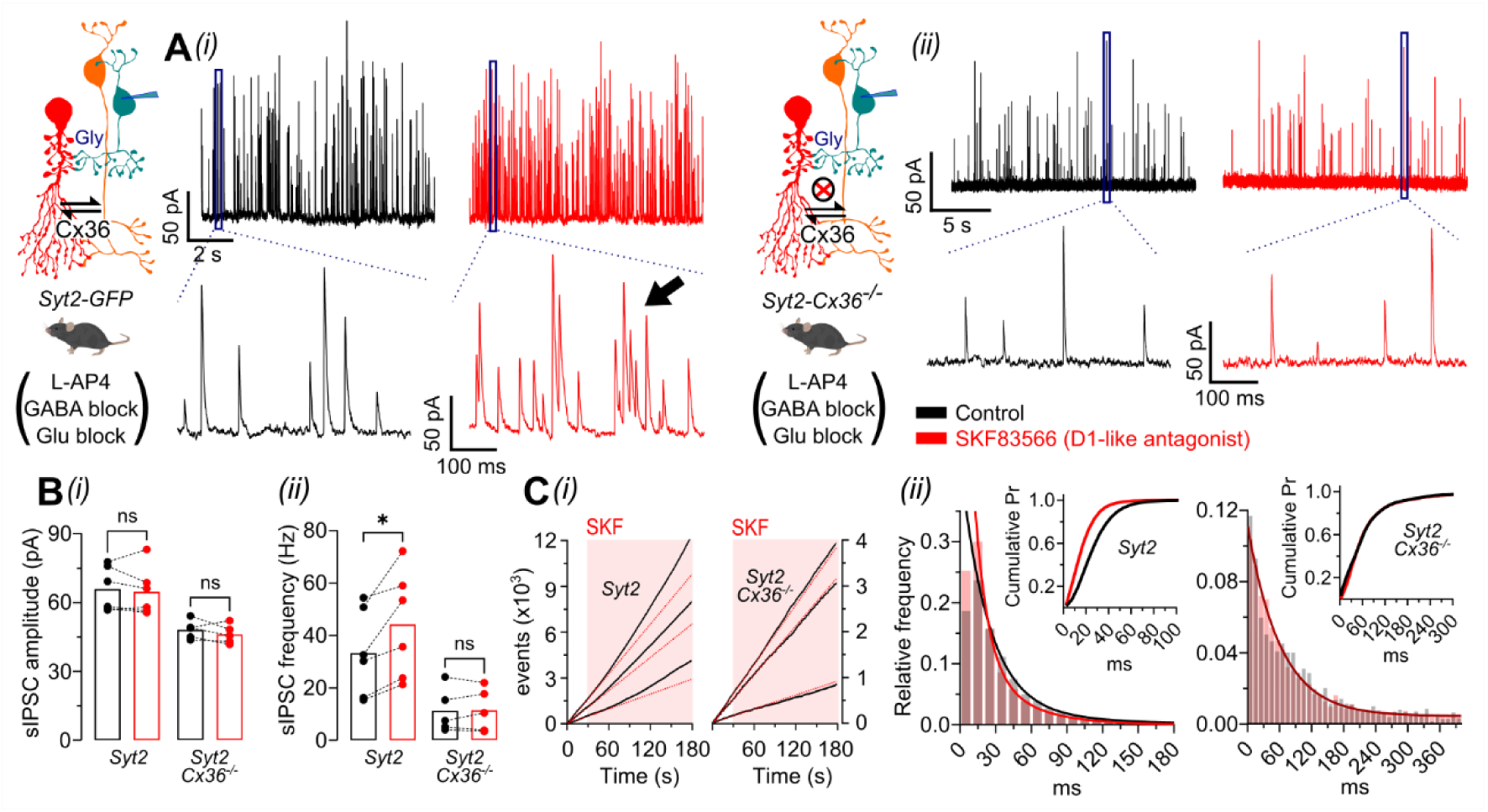
Blocking D1Rs enhanced AII-AC glycinergic transmission to OFF-CBCs in *Syt2* but not *Syt2-Cx36^−/−^* animals. **(A)** Example voltage-clamp recordings of glycinergic sIPSCs (holding = 0 mV) in type-2 OFF-CBCs from *Syt2-GFP **(i)*** and *Syt2-Cx36^−/−^ **(ii)*** animals, before and after application of the D1R antagonist SKF83566. Note the absence of effect in *Syt2-Cx36^−/−^* animals, a result consistent with the findings in pre-synaptic AII-ACs. Overlapping sIPSCs (inset *black arrow*) were typically observed after SKF83566 application in *Syt2-GFP* animals. Note also that both frequency and amplitude of sIPSCs are smaller in *Syt2-Cx36^−/−^* animals. To isolate the spontaneous glycinergic events, a synaptic cocktail was added to the external solution. **(B)** Mean sIPSC amplitude (only isolated, non-overlapping events) ***(i)*** and mean sIPSC frequency ***(ii)*** before and after SKF83566 for each mice strain. **(C)*(i)*** Examples of sIPSC time series (3 example cells over 3 minutes) for OFF-CBCs from each mice strain. Note that in *Syt2-GFP* animals SKF83566 slowly increases sIPSC frequency. Red dotted lines correspond to a linear regression over the first 30 seconds before the drug reaches the bath. ***(ii)*** sIPSCs inter-event interval (IEI) distributions, along with cumulative probability plot (*inset*), in control and after SKF83566, for each mice strain. IEI distributions were fitted by different double exponentials in *Syt2-GFP* animals, and a single exponential (*dark red*) fitted both distributions (control and treatment) in *Syt2-Cx36^−/−^* animals. *Syt2-GFP*: *n* = 6 cells; Ctrl: 4727 ± 2680 events per cell, SKF83566: 6675 ± 4550. *Syt2-Cx36^−/−^*: *n* = 5; Ctrl: 949 ± 772 events per cell, SKF83566: 1686 ± 1294 events per cell. Circles in bar graphs represent single cells. Statistical significance was determined using paired *t*-tests; ns: *p* > 0.05; *: *p* < 0.05.

In conclusion, our results strongly support the idea that modulation of AII-AC’s Cx36 gap junctions by D1Rs plays a key role in the spontaneous or tonic glycinergic transmission to OFF-CBCs. Combining the data presented here with that from the previous experiment (**Figure 5**), we suggest that even if D1R-mediated changes in gap junctions result in only small alterations in the V_m_, the impact on the spontaneous glycine release can still be quite strong.

### Synaptic blocker cocktail alters resting V_m_ and input resistance in AII-ACs

In the previous sections, we used a synaptic blocker cocktail, consisting of the AMPA/kainite receptor blocker CNQX, the NMDA receptor blocker DL-AP5, the GABA_(A)_ receptor blocker gabazine, and the group-III mGluR agonist L-AP4, to isolate glycinergic synapses between AII-ACs and OFF-CBCs and permit the analysis of dopaminergic modulation of AII-AC signaling. However, this cocktail of synaptic blockers led to a retinal state where D1Rs in AII-ACs appear to be saturated (**Figure 4**). Therefore, it was important to examine the effects of the synaptic blocker cocktail on AII-AC resting V_m_ and R_input_.

Using a current step protocol in whole-cell current-clamp, we found that application of the synaptic blocker cocktail significantly altered the membrane voltage responses to current injections (**Figure 7A**). Compared to before the drug wash (**Figure 7A*(i)***), AII-ACs post-cocktail displayed larger voltage responses to each current injection (**Figure 7A*(ii)***), suggesting a higher input resistance in AII-ACs after the cocktail. Additionally, since both excitatory and inhibitory inputs from other neurons were blocked, baseline membrane voltages fluctuations were reduced, and spikelets were easily distinguished from background synaptic noise. When expanded, the dramatic differences in AII-AC V_m_ stability before (**Figure B*(i)***) and after (**Figure B*(ii)***) the synaptic blocker cocktail become easily visible. The cocktail also reduced the AII-AC resting V_m_ from −32 ± 4.5 mV to −40.2 ± 2.*6* mV, *p* = 0.0033, and increased R_input_ from 332 ± 67 MW to 378 ± 62 MW, *p* = 0.0088. These results indicate that the cocktail dramatically changed the electrophysiological properties of the AII-ACs and likely that of all retinal neurons, including DA-ACs and their tonic release of dopamine.

**Figure 7.**
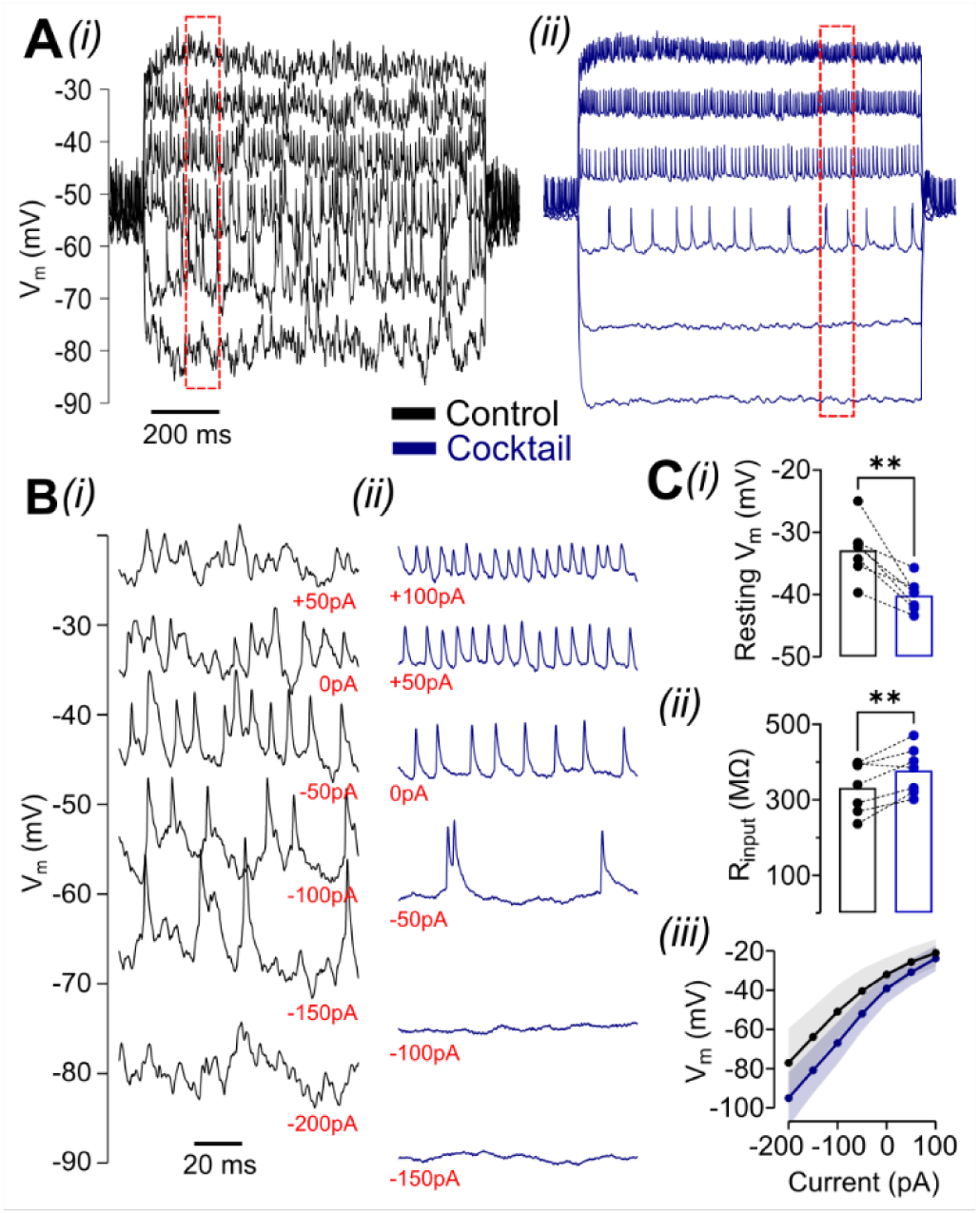
Synaptic blocker cocktail altered excitability and input resistance of AII-ACs. **(A)** Current-clamp recording from an AII-AC after 1 second current steps before ***(i)*** and after ***(ii)*** the application of a synaptic blocker cocktail containing the AMPAR blocker CNQX (10µM), the NMDAR blocker DL-AP5 (50µM), the GABA(A)R blocker gabazine (8µM) and the group-III mGluR agonist L-AP4 (10µM). **(B)** Voltage traces at several current injections show dramatic differences in AII-AC Vm and spiking before ***(i)*** and after ***(ii)*** the cocktail. Current step injection values are shown in red. **(C)** Quantification of resting membrane voltage ***(i)***, input resistance at −55 mV ***(ii)***, and IV curve ***(iii)*** in control (*black*) and cocktail (*blue*). *n* = 7. *Circles* in bar graphs represent single cells. Statistical significance was determined using paired *t*-tests; **: *p* < 0.01.

## Discussion

We have investigated how dopamine modulates the V_m_ and spiking properties of AII-ACs in both *wt* and *Cx36^−/−^* mice. Using the *Syt2-GFP* and the *Syt2-Cx36^−/−^* transgenic mice, we demonstrated that AII-AC resting V_m_, input resistance, spiking properties, and glycine transmission to downstream OFF-CBCs, are controlled by D1R-mediated changes in gap junctions between AII-ACs and ON-CBCs. We note that single-cell RNA transcriptomics data shows moderate expression of D1Rs in type-6 ON-CBCs (Shekhar et al., 2016), which are thought to be the ON-CBC type most likely to be coupled to AII-ACs (Graydon et al., 2018), so we cannot rule out that D1R modulation of AII-AC properties and synaptic output could be bolstered by dopamine regulation of ON-CBCs (Farshi, Fyk-Kolodziej, Krolewski, Walker, & Ichinose, 2016; Hellmer, Bohl, Hall, Koehler, & Ichinose, 2020).

### The relationship between V_m_, spiking properties and glycinergic transmission of AII-ACs

Spikelets in AII-ACs are generated by fast Na channels (Cembrowski et al., 2012; Wu, Ivanova, Cui, Lu, & Pan, 2011), which have been shown to be subject of dopaminergic modulation in RGCs (Hayashida et al., 2009; Ichinose & Lukasiewicz, 2007). In order to interpret how dopamine and the D1Rs agonists/antagonists could affect AII-ACs excitability and firing, we first investigated the extent by which the spiking properties would vary with V_m_ in control conditions. We found a strong relationship between AII-AC V_m_ and spiking properties (**Figure 1 and Figure 7**). At hyperpolarized potentials, below −70 mV, intrinsic spiking was absent and EPSPs were not able to elicit spiking behavior. At around −65 mV, we began to see occasional spikes, but bursting predominated. At even more depolarized potentials, such as −45 to −40 mV, burst spiking had been completely replaced by high-frequency, low-amplitude spikelets that were eventually abolished as V_m_ approached −25 mV and Na^+^ channel inactivation became very significant.

The relationship between V_m_ and spiking properties of AII-ACs has been previously studied, although burst analysis was not attempted (Tamalu & Watanabe, 2007). In agreement with our results, the more depolarized the AII-ACs, especially between −65 and −40 mV, the higher the overall spike frequency. We observed the opposite for bursts, and therefore, it’s possible that low-voltage or subthreshold ion channels are critical to generate them. Tamalu & Watanabe (2007) also showed that application of L-AP4 hyperpolarized AII-ACs and inhibited their spiking. It was therefore suggested that the inhibition of RBCs by L-AP4 leads to reduced glutamatergic input to AII-ACs. However, by knocking out gap junctions using the *Cx36^−/−^* mouse, we found that electrical synapses between AII-ACs and ON-CBCs play a key role in setting the resting V_m_, suggesting that intrinsic inward currents and excitatory inputs from upstream ON-CBCs may both help to control AII-AC spiking. We also found that the frequency of sIPSCs in type-2 OFF-CBCs is directly correlated with AII-AC resting V_m_ (**Figure 6**). These complementary results suggest that AII-ACs convert both rod and cone ON responses into spike frequency and V_m_ depolarization, most likely in a light intensity-dependent manner. Small inputs from either RBCs or ON-CBCs at low light intensity elicit low-frequency spikes and reduced glycinergic inhibition to OFF-CBCs and OFF-RGCs. Larger inputs from upstream neurons at higher intensity light will elicit high-frequency spiking behavior and high rates of glycinergic inhibition to downstream neurons.

### AII-ACs respond to D1R agonists differentially in the presence or absence of synaptic blockers and L-AP4

AMPA/kainate receptors are expressed throughout the retina, including OFF-bipolar cells (DeVries, 2000), AII-ACs (Singer & Diamond, 2003), and DA-ACs (Liu, Alessio, Spix, & Zhang, 2019). NMDA receptors are also present in AII-ACs (Margaret L. Veruki, Zhou, Castilho, Morgans, & Hartveit, 2019) and DA-ACs (Liu, Spix, & Zhang, 2017), while GABA_(A)_ receptors are found in many retinal cell types, including bipolar cells (Fletcher, Koulen, & Wässle, 1998; Schubert, Hoon, Euler, Lukasiewicz, & Wong, 2013) and AII-ACs (Beltrán-Matas, Hartveit, & Veruki, 2023). Indeed, there’s some evidence that DA-ACs themselves may develop conventional GABAergic synapses directly onto AII-ACs (Contini & Raviola, 2003). mGluR6, a group-III mGlu receptor unique to the retina, is the primary glutamate sensor in ON bipolar cells (Vardi & Morigiwa, 1997) that controls the RBC and ON-CBC light response. Accordingly, it is reasonable to predict that pharmacological blockade of these widely-expressed receptors may significantly alter electrophysiological properties throughout the retina, including in AII-ACs and DA-ACs. Retinal DA-ACs are known to exhibit an intrinsic or pacemaker firing, which enables them to spontaneously release large amounts of dopamine. This spontaneous activity is then strongly modulated by the balance between excitatory and inhibitory inputs received from ON-CBCs and GABAergic ACs, respectively, with glycinergic ACs possibly playing a less significant role (Feigenspan, Gustincich, Bean, & Raviola, 1998; Marshak, 2001; Newkirk, Hoon, Wong, & Detwiler, 2013; Pérez-Fernández et al., 2019; Puopolo et al., 2001; Steffen et al., 2003). In the model proposed by Marshak (2001), ON-CBCs and GABAergic ACs would be hyperpolarized in the presence of the mGluR6 agonist APB and a photopic background. Synaptic input to DA-ACs would be blunted, causing them to be spontaneously active. With the synaptic blocker cocktail and L-AP4, DA-ACs would now be devoid of synaptic input and could even exhibit a higher level of spontaneous activity, perhaps with a more depolarized V_m_ and higher spike frequency. Therefore, it is reasonable to expect that retinal dopamine levels may be strongly augmented by the drug cocktail, resulting in an artificial condition where D1Rs are fully saturated. Indeed, we found that use of the cocktail eliminated the effect of the D1R agonist on AII-AC membrane voltage and prevented subsequent changes in postsynaptic OFF-CBCs (**Figure 4**). However, D1R antagonists still had an effect, suggesting that D1Rs in AII-ACs may be saturated (**Figure 5 and Figure 6**). This saturation did not occur consistently in the absence of the synaptic blockers (**Figure 2 and Figure 3**) but was seen in about half of AII-ACs. In the absence of the cocktail and in dim photopic conditions, we propose that some AII-ACs may still have their D1Rs saturated because dopamine levels may be high in some retinal slices due to outcompeting excitatory inputs to DA-ACs. As a consequence of slice preparation, both the balance between inhibitory and excitatory inputs, as well as the overall health of DA-ACs can be disrupted. Additionally, AII-ACs may be heterogenous in their expression of D1Rs and some AII-ACs may show absent or very low levels of D1R expression, as suggested by single-cell RNA-seq data (Yan et al., 2020).

### Dopamine modulates AII-AC membrane potential and transmission to OFF-CBCs by regulating gap junction coupling with ON-CBCs

Exogenous dopamine and application of a D1R agonist both hyperpolarized the AII-AC resting V_m_, while a D1R antagonist depolarized it (**Figures 2 and 3**). The latter effect was absent in *Cx36^−/−^* retinas (**Figure 5**). Furthermore, blocking D1Rs caused an increase in the frequency of sIPSCs in postsynaptic OFF-CBCs, an effect that was also absent in *Cx36^−/−^* retinas (**Figures 4-6**). The correlation between the D1R antagonist depolarization and the change in R_input_ of AII-ACs, the transmission of an inward current to AII-ACs and the strong reduction in sIPSC frequency in OFF-CBCs from Syt2-Cx36^−/−^ animals (**Figure 6Bii**) all point towards modulation of gap junctions between AII-ACs and ON-CBCs. We believe these results clearly demonstrate a major mechanism by which D1Rs affect AII-AC glycinergic transmission to OFF-CBCs. Mazade et al. (2019) have previously shown that D1 receptor activation reduced light-evoked and spontaneous inhibition of OFF-CBCs to light-adapted levels. The authors pointed to reduced glycine release by AII-ACs as the most likely causal explanation for their results, and here we have not only provided further support to this hypothesis but also revealed the presynaptic mechanisms involved. Early studies showing that D1R antagonists lead to hyperpolarization and reduced spontaneous activity in OFF-RGCs, presumably due to increased glycinergic inhibition, also support our findings (Jensen, 1992; Jensen & Daw, 1986).

The role of gap junction communication between AII-ACs and ON-CBCs in amplifying crossover inhibition has been recently demonstrated (Shubhash Chandra Yadav, Ganzen, Nawy, & Kramer, 2024). By optogenetically depolarizing type-6 ON-CBCs, the authors were able to simultaneously record robust IPSCs in OFF-sustained RGCs. The IPSCs were blocked by either MFA or strychnine. These findings agree with our experiments showing that D1R antagonist depolarization of AII-ACs increases glycinergic sIPSCs in type-2 OFF-CBCs. It’s interesting that Yadav et al. performed their experiments under photopic conditions, which suggests a substantial gap junction conductance between AII-ACs and ON-CBCs even when dopamine levels are presumably high. Importantly, we observed a much higher R_input_ in *Cx36^−/−^* AII-ACs than in *wt* AII-ACs under the drug cocktail when D1Rs are apparently saturated by endogenous dopamine. Therefore, it’s reasonable to suppose that blockade or activation of D1Rs modulates the gap junction conductance only to a limited, but physiologically relevant, extent.

Another important finding was that the D1R antagonist decreased the intrinsic excitability of AII-ACs: spikelet frequency measured at the same V_m_ was lower after drug application. This is probably a consequence of the decrease in R_input_ we observed in the experiments with the drug cocktail, a result consistent with enhanced coupling to ON-CBCs and others AII-ACs. We also found evidence that the D1R agonist causes the opposite effect in the absence of the cocktail. We haven’t measured the R_input_ in this particular experiment, but a prediction is that it will increase following D1R activation. Yadav et al. (2024) reported that the D1R antagonist SCH23390 strongly depressed the synaptic output of ON-CBCs in a TTX-independent manner, a result that could be explained by decreased R_input_ in both AII-ACs and ON-CBCs. The authors suggest that the Na^+^ currents present only in AII-ACs would be shunted by the increased coupling between AII-ACs themselves. Reduced excitability in ON-CBCs was also directly demonstrated by paired optogenetic stimulation and EPSC recording in ON-RGCs, a result that is consistent with our findings in AII-ACs.

Interestingly, application of the D1R agonist did not completely recapitulate the effects of exogenous dopamine on AII-AC activity (**Figure 2**). While both dopamine and the D1R agonist resulted in significant AII-AC hyperpolarization, the effect of dopamine was faster, causing a noticeable V_m_ hyperpolarization within approximately 150 seconds compared to 250 seconds for the D1R agonist (**Figure 2**). Additionally, dopamine-mediated hyperpolarization was accompanied by a marked spike frequency reduction. The slow onset of hyperpolarization after the D1R agonist is consistent with the slow modulation of gap junction coupling via dephosphorylation of Cx36 (Kothmann et al., 2009). The fact that exogenous dopamine exhibits an effect on faster time-scales suggests that additional mechanisms may be at play, likely circuit-level mechanisms involving synaptic release and the activation of D1Rs and D2Rs in other cell types.

### Implications for visual processing

Our findings about D1R modulation of spontaneous glycinergic transmission from AII-ACs can offer insights into how the inner retina circuits adapt to slow changes in environmental light levels (e.g. light adaptation to sustained light stimuli). Our results suggest that, even in the condition where excitatory input is blocked (and ON-BCs are hyperpolarized by L-AP4), AII-ACs still provide strong tonic inhibition to OFF-CBCs, which certainly impacts how these cells respond to light (Graydon et al., 2018). We often observed very high frequency glycinergic sIPSCs (>40 hz) under these conditions. Intermediate levels of dopamine release, as expected in low photopic conditions, could potentially saturate a fraction of D1Rs, which in turn would hyperpolarize AII-ACs, and simultaneously contribute to an increase in their excitability through gap junction uncoupling and increased R_input_. The combination of these effects can support efficient coding of EPSCs from RBCs and reduce synaptic noise between AII-ACs and OFF-CBCs. Under conditions where dopamine levels are low, as in dim light, AII-ACs are highly coupled, less excitable, and have large receptive fields. Inactive D1Rs could then contribute to the depolarization of AII-ACs cause by excitatory inputs from RBCs and inward currents from ON-CBCs through gap junctions, and consequently, tonic inhibition to OFF-CBCs would be strong. Visual stimuli causing a decrease in the frequency of sIPSCs in the OFF-CBCs, such as a dark object in twilight, would be coded more efficiently, and discrimination and contrast of these objects could be improved.

High levels of dopamine during bright light conditions would activate D1Rs, which would strongly hyperpolarize AII-ACs and inhibit their spiking. Tonic inhibition of OFF-CBCs would be diminished, allowing ON signals from ON-CBCs to be faithfully transmitted to OFF-CBCs, via crossover inhibition, as an increase in sIPSC frequency. This form of processing may improve contrast sensitivity in bright backgrounds. Moreover, hyperpolarization of AII-ACs could prevent or reduce transmission of noise from saturated RBCs to ON-CBCs, and consequently ON-RGCs. This signal-to-noise enhancing mechanism could be important during light-adaptation in the mammalian retina, especially because saturated rod photoreceptors utilize the cone pathway but provide little useful information for vision under high photopic conditions (Roy & Field, 2019). Overall, these dopaminergic modulatory mechanisms appear critical to fine-tune the levels of tonic inhibition received by OFF-CBCs and the activity of ON-CBCs, which in turn could shape how these cells respond to changes in environmental light and diverse light stimuli. These mechanisms may help optimize spike encoding in OFF-RGCs to improve contrast detection under different light conditions.

## Author contributions

Conceptualization: PSJ, CMW, HvG; Data curation: PSJ, CMW; Formal analysis: PSJ, CMW; Funding acquisition: CMW, HvG; Investigation: PSJ, CMW; Methodology: PSJ, CMW, HvG; Project administration: HvG; Software: PSJ, CMW; Resources: PSJ, CMW, HvG; Supervision: HvG; Validation: PSJ, CMW, HvG; Visualization: PSJ, CMW, HvG; Writing – original draft: PSJ, CMW, HvG; Writing – review and editing: PSJ, CMW, HvG

